# Hydrogen properties and charges in a sub-1.2 Å resolution cryo-EM structure revealed by a cold field emission beam

**DOI:** 10.1101/2021.12.21.473430

**Authors:** Saori Maki-Yonekura, Keisuke Kawakami, Tasuku Hamaguchi, Kiyofumi Takaba, Koji Yonekura

## Abstract

The cold field emission (CFE) beam produces the less-attenuated contrast transfer function of electron microscopy, thereby enhancing high-resolution signals and this particularly benefits higher-resolution single particle cryogenic electron microscopy. Here, we present a sub-1.2 Å resolution structure of a standard protein sample, apoferritin. Image data were collected with the CFE beam in a high-throughput scheme while minimizing beam tilt deviations from the coma-free axis. A difference map reveals positive densities for most hydrogen atoms in the core region of the protein complex including those in water molecules, while negative densities around acidic amino-acid side chains likely represent negative charges. The position of the hydrogen densities depends on parent bonded-atom type, which is validated by an estimated level of coordinate errors.

---

Excellent coherence of an electron beam is now recognized as an important factor for high-resolution single-particle cryogenic electron microscopy (cryo-EM) (1, 2, 3, 4). The cold field emission (CFE) gun produces a high temporal-coherence beam under high vacuum without the application of heat. The energy spread of the CFE beam is less than half of that of standard thermal field emission or Schottky emission (SE), which requires heating of the cathode tip to a few thousand degrees Kelvin (K). The contrast transfer function (CTF) for the CFE source is remarkably less attenuated at a resolution range beyond 2 Å than that from the SE source at the same defocus value (1). A monochromator can produce an even smaller energy spread (2), but the device may prove impractical for many users due to instability, additional complexities in electron optics and operation, and high price. High-resolution signals are clearly visible in Fourier-transforms of metal images recorded with the CFE electron beam (Fig. 1A and Figs. 2 and 3 of Ref. 1).

**Figure 1.**
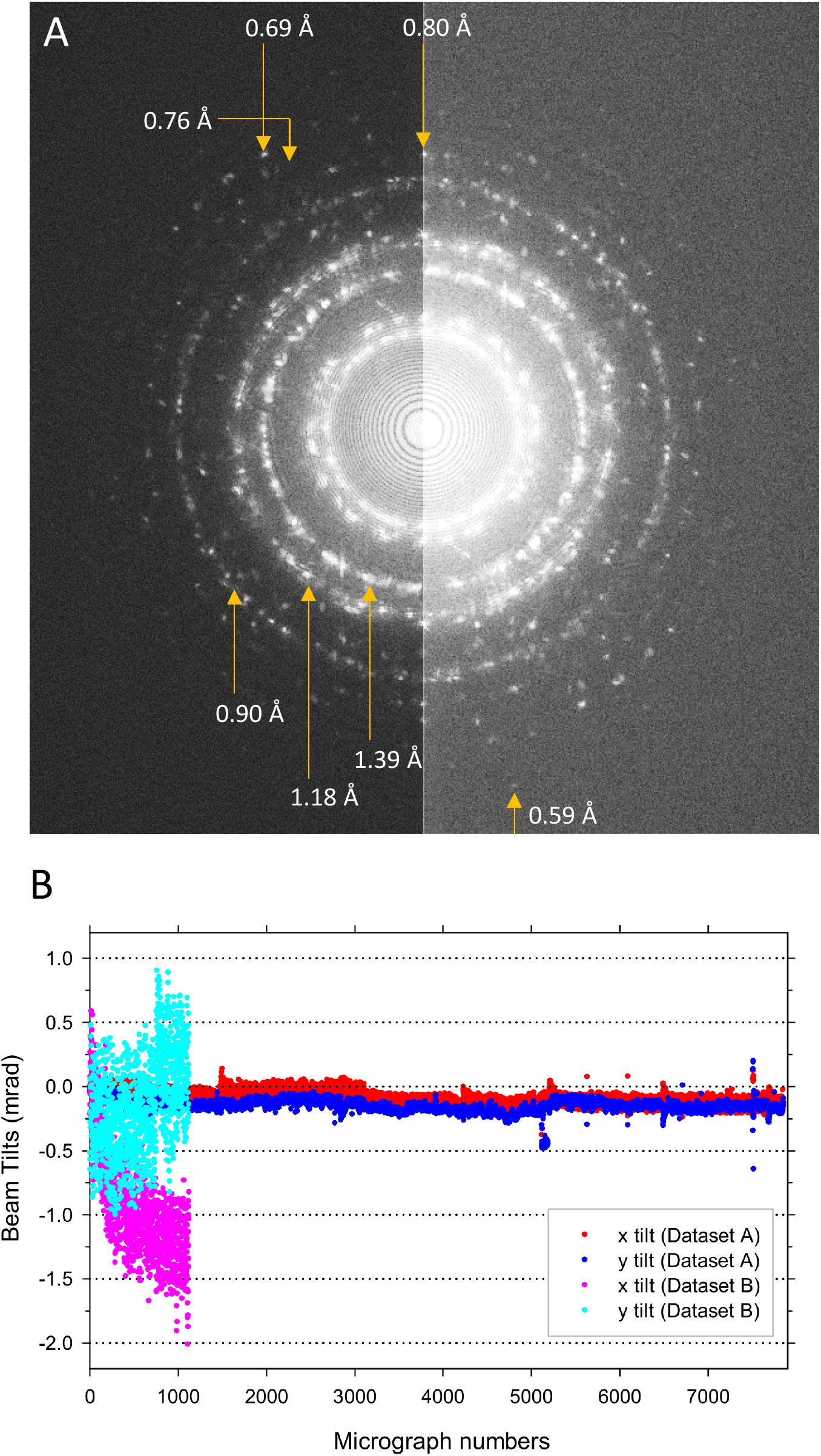
Characteristics of the electron beam and datasets for cryo-EM analyses. (A) Fourier transform of a metal image. Carbon film deposited with Pt-Ir was imaged with a CFE gun at a nominal magnification of × 300,000 and defocus of ~ −1 μm. Beam illumination was focused and marginally off the parallel condition in order to give significant counts in each frame. Movie images were recorded on a K3 detector in counted mode at a dose rate of ~ 7 e- / pixel / s for 15 s exposure, and motion-corrected full frames were summed with DigitalMicrograph. Signals to ~ 0.6 Å are visible. The right half is brightened for clarity of high-resolution signals. (B) Beam tilt variations during data collections. Compensation of beam shifts were preadjusted in imaging mode (Dataset A) and in diffraction mode (Dataset B).

**Figure 2.**
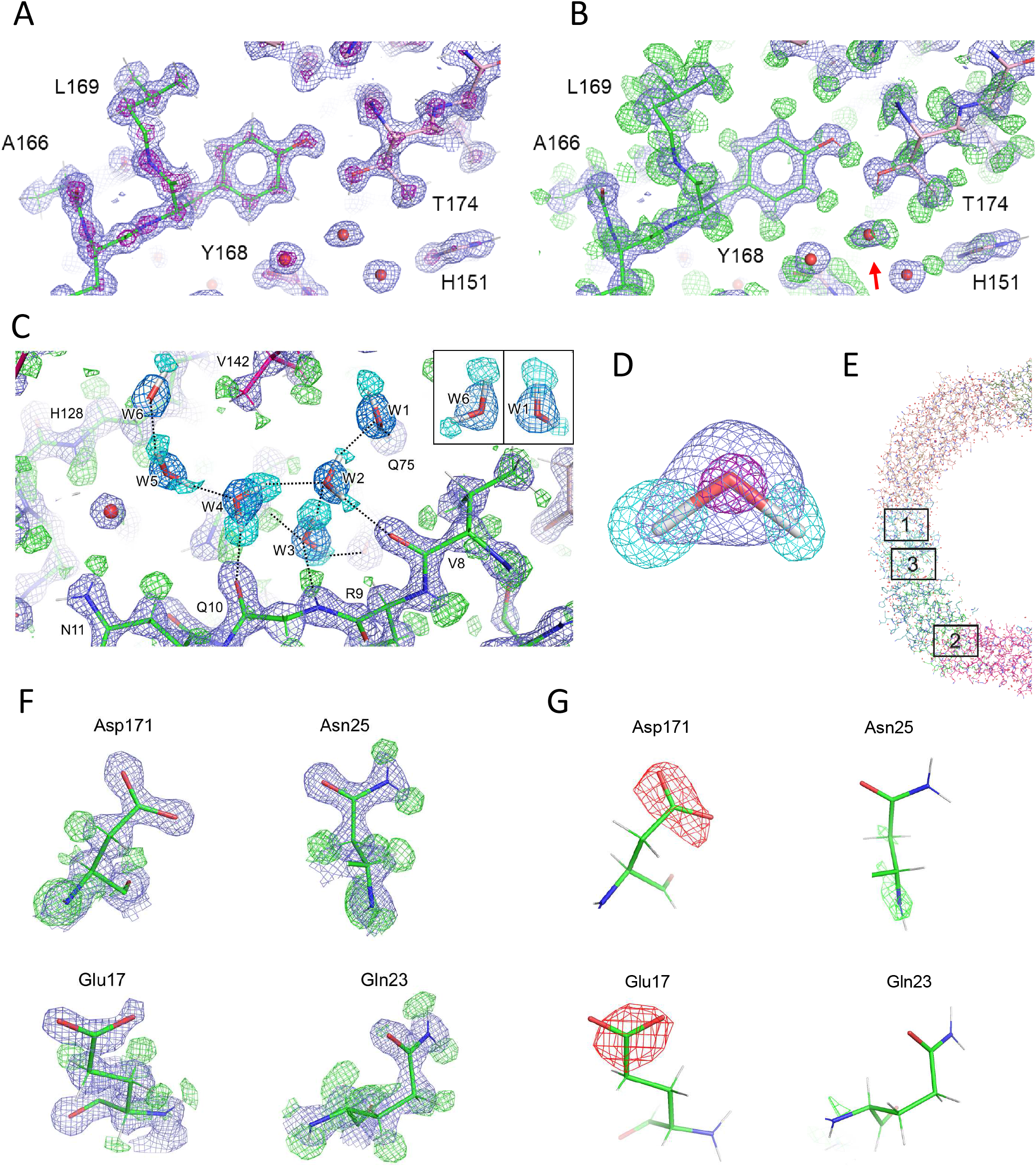
Structure details of apoferritin. (A, B) Around a tyrosine (Tyr 168) overlaid with the atomic model refined in this study. Experimental densities are shown in blue (A, B) and purple (A) and positive densities in a difference map between the experimental data and hydrogen-omitting model in green (B). Arrow in B indicates a water molecule with two hydrogen densities in green. (C) Around a water cluster. Sky blue nets represent six water molecules (W1 – W6) showing two hydrogen densities in cyan and comprising a water cluster through hydrogen bonds. (D) One of the water molecules (W4) in C. (E) A cutout view of half the apoferritin complex indicating the area marked “1” for A and B, “2” for C and “3” for Figs. S2A and B. (F) Zoom-up views of single amino-acid residues, aspartate (Asp 171), asparagine (Asn 25), glutamate (Glu 17), and glutamine (Gln 23) with experimental densities in blue and positive densities in the difference map in green. (G) The same as in F, but with positive and negative densities in green and red, respectively, in the difference map calculated by limiting the data to 2.5 Å resolution. Blue nets in A–D and F and purple nets in A are displayed at density levels of 2*σ* and 7*σ*, respectively. Sky blue nets for water in C is at 2*σ*. Green nets in A-D and cyan nets in C and D are at 2.5 - 3*σ*. Cyan nets in insets of C are at 2*σ* for clarity of W1 and W6. Green nets in F show differences at 3*σ*, and green and red nets in G are 4 and −4*σ*, respectively.

Imaging with the improved CTF by the CFE beam or through the monochromator has yielded single particle reconstructions of the standard protein complex apoferritin at 1.22 Å and 1.25 Å resolutions (3, 2). These analyses even show hydrogen densities in difference maps (3, 5). Hydrogen and hydrogen bonding play critical roles in stabilization of protein structures and functions such as enzymatic catalysis, electron transfer and ligand binding (e.g. 6, 7, 8), but identification of hydrogen atoms is often hampered by a low visibility for X-rays (e.g. 8, 5).

Our previous data taken using a CFE gun equipped in a CRYO ARM 300 microscope (JEOL) contained large fluctuations in beam tilts, even though such images were collected through mechanical stage shifts exclusively without introducing additional beam tilts (1, 9, 4, 10, 11). The data gave reconstructions to ~1.5 Å resolution (1, 4). Beam tilts from the axial coma-free axis cause a marked increase in phase errors in proportion to the third power of spatial frequencies. Although it is possible to estimate the beam tilts in acquired images and correct errors (12, 13, 14), such a post-correction scheme has its limitations particularly for larger beam-tilt errors at higher resolutions.

Here, we report a sub-1.2 Å resolution structure of apoferritin reconstructed from images collected rapidly using image and stage shifts for positioning data-taking areas while keeping the magnitude of beam tilt deviations small (12). The cryo-EM map resolves separate densities for individual atoms, many hydrogen atoms and multi-conformations of amino acid residues. We then analyse the position of the hydrogen atoms and its dependence on bond type. We also show that appearance of negative densities in a difference map varies depending on spatial frequencies, which likely represents negative charges. Of course, charges are critically important for macromolecules functioning as well. Subatomic resolution X-ray crystallography provides charge distributions inside proteins experimentally (15, 16, 17), but such studies are limited.

## Results and Discussion

### Minimization of beam tilts off the axial coma-free axis

Large fluctuations of beam tilts have been found in previous data collections, sometimes ranging over several mrad (1, 9, 4, 10, 11), and we suspected that this is due to off-pivot points of the beam on the specimen plane. Adjustment of the pivot points needs accurate compensation for beam tilts and shifts by means of condenser lens deflector coils. We had used the diffraction mode for compensation of beam shifts, as per the manual provided by JEOL. Beam tilt fluctuations still appeared significant and were not improved by simply changing the magnification without applying any image shifts. The fluctuations became much worse during data acquisition in a high throughput scheme where both stage and image shifts were implemented and one image stack per hole and 3 × 3 stacks from holes clustered around a central hole were taken once the stage is moved to a new position (Figs. 1B and S1B). This happened even though we had used active adjustment of beam tilt to the axial coma-free axis based on calibration by SerialEM (18) during data collection. Post-estimation and correction of beam tilts for all image stacks separately provided Dataset B, which gave a 3D reconstruction to ~1.49 Å resolution by the gold standard Fourier shell correlation (FSC) criterion (Figs. S1C and F) (19).

We then adjusted compensation of beam shifts in imaging mode as was done for the data acquisition condition (see Materials & Methods and ref. 20). SerialEM and JAFIS Tool (see Method) were used to take image stacks over 5 × 5 holes per stage shift. This resulted in remarkably improved beam tilts in the individual images comprising this dataset, which we named Dataset A, and most of them show deviations within ~ 0.15 mrad (Figs. 1B and S1A), even though larger image shifts were applied during image acquisition than had been used for Dataset B. The obvious improvements provided by the new alignment method suggests that the effective sample height changes between diffraction and imaging modes. Thus, for the CRYO ARM microscope, it is important to adjust compensation of beam shifts under the condition as used for data acquisition.

More than 2,000,000 particles from ~ 7900 image stacks gave a 1.21 Å resolution map, and the final resolution reached 1.19 Å (Figs. S1C and E) by rescaling to a finer sampling over super-resolution pixels of the K3 detector. The estimated Rosenthal-Henderson B-factor (Fig. S1D) (21) is comparable to the previous reconstructions (2, 3). The Rosenthal-Henderson plots for Datasets A and B appear similar up to ~ 1.7 Å, with the most prominent difference being the beam tilt fluctuations (Figs. 1B, S1A and B). Thus, post-correction of beam tilts is probably effective to this resolution range, but becomes less so beyond that with large beam tilt fluctuations.

The catalogue value of the spherical aberration coefficient, *Cs* = 2.7 mm, means that even 0.1 mrad off-estimation of beam tilt introduces significant phase errors in high-resolution ranges (Fig. S1G). Errors are much worse with an accelerating voltage of 200 kV compared with 300 kV, and a new corrector was introduced to remove phase errors by both on-axial and off-axial coma (2). Nevertheless, our data indicate that the resolution can be extended to sub-1.2 Å without this device. Again, a corrector would be impractical for the same reasons as for the monochromator. We used simple holey carbon film on copper grids for sample support, but use of gold grids may further improve the resolution (22).

### Structure features

The cryo-EM map resolves separate densities for individual atoms particularly in the core region of the protein complex (Figs. 2 and S2A), while side chains at the surface of the apoferritin spherical shell are rather flexible and some residues show multi-conformations (Fig. S2C). These features were also observed in previous structure analyses (2, 3). The high quality of the cryo-EM map allowed us to build the atomic model unambiguously, and the model metrics are good (Table S1).

Then, we calculated a difference map between the experimental data and the model omitting hydrogen atoms by *Servalcat* (5). The difference map reveals many densities corresponding to hydrogen atoms. The number of hydrogen atoms associated with the protein model is 905, which corresponds to ~ 70% of total possible hydrogen atoms in the model, after peaks at 2σ were edited by visual inspection. In the protein core, most of the hydrogen atoms are identified for both main chain and side chains. Representative structures are shown in Figs. 2, S2B, and S2D. Structures around a tyrosine (Tyr 168) and a phenylalanine (Phe 51) clearly exhibit all the hydrogen atoms. The hydrogen density in the hydroxyl group of Tyr 168 is bent towards an oxygen atom in a neighbouring residue for hydrogen bonding (Figs. 2B and S2D). Some water molecules also show resolved hydrogen densities, and we identified six water molecules with two separate hydrogen densities, which contribute to formation of a water cluster through hydrogen bonding (Fig. 2C). One of them, shown in Fig. 2D, fits well to the configuration of a theoretical water model.

Typical maps are presented for several amino acid types in Fig. 2F. Reflecting the pH 7.5 of the sample solution, acidic amino acid residues appear to be de-protonated. The amide and oxygen termini in asparagine and glutamine can be clearly identified by the presence or absence of hydrogen densities (Fig. 2F). Oxygen termini in aspartate and glutamate are also well resolved, although the side chains of aspartate and glutamate are known to be particularly susceptible to radiation and sometimes disappear by exposure to a high X-ray dose (23). In the protein structures obtained by electron 3D crystallography (3D ED), we observed decreased densities for negatively charged atoms (24), since electron scattering factors of anions become negative or nearly zero at lower spatial frequencies (24, 25, 26). A cut of structure factors in lower resolutions recovered some densities for the side chain of aspartate residues (21). In single particle analysis, amplitude information is less accurate due to the effects of CTF and the standard post-process procedure sharpens the map. This amplification of high-resolution parts and/or amplitude modification appear to maintain densities of acidic amino acid side chains in the experimental maps (Fig. 2F). While, in the difference map subtracting structure factors consisting of neutral atoms from the experimental data, there are a negative density at a density level ≤ −4*σ* near the terminal oxygen atom of Gln 23 but larger and more dispersed negative densities at the same level around the side chains of Asp 171 and Glu 17 (Fig. S3A). Here, we selected these residues with estimated temperature factors (B-factors) of terminal oxygen atoms < 25 Å^2^ except for Glu 17. Removal of the data in higher than 2.5 Å reveals more prominent negative densities only on the acidic side chains (Figs. 2G and S3B). Such negative densities decrease gradually upon cut of lower-resolution terms (Fig. S3C). These observations should reflect the fact that the electron scattering factors vary considerably between the neutral and charged atoms but match from ~ 2.5 Å to higher resolutions (24, 25, 26). It is obviously contradictory to radiation damage, as radiation damage must be more severe for higher-resolution structures (cf. Figs. S3B and C). Thus, the observed negative densities in Figs. 2G and S3 would rather reflect the charged state of the amino acids.

The analysis above is still qualitative. Difference maps could provide smaller differences between the experimental data and model, but our recent 3D ED study showed that interpretation of residual densities in a difference map is more challenging due to distributions of partial charges and suboptimal assignment of electron scattering factors to the corresponding model (27). Conversion to electron density may provide a more interpretable result (27). Still, further work is needed for a more extensive analysis of charges.

### Hydrogen properties

In single particle reconstructions and Coulomb potential maps by 3D ED, hydrogen densities appear further away from their parent atoms than those in X-ray crystal structures (28, 3, 27). This reflects the fact that incident electrons are affected by both electrons and nuclear charges and the latter is dominant, while X-rays are scattered by electrons around atoms and the electron in a hydrogen atom is attracted towards its parent bonded atom (29, 30). The electron beam makes the hydrogen densities appear even slightly further from the bonded atom than the nucleus of hydrogen (3, 27). We have shown that in the Coulomb potential map of an organic molecule the hydrogen density peak position depends on the type of covalent bond (27). There is a larger change in the position of the hydrogen peaks in electron density maps depending on the polarity of bonding due to the mobility of electrons (29, 30, 31). In general, the higher the attractive force shortens the bond length (29, 27), while conversely higher B-factors and lower resolution lengthen the apparent distance (3).

We summarize the statistics on distances of hydrogen peaks from parent atoms in the 1.19 Å cryo-EM map in Table 1. The averaged distances at with density levels ≥ 4*σ* are almost identical to the nuclear positions for all the bond types except for O-H bonds. The number for O-H bonds is probably too small for reliable estimation of the distance. Hydrogen peaks in polar bonds (N-H) appear shorter than those in other C-H_n_ bonds. This matches well with the results obtained by 3D ED (27). For X-rays, the peak distance in the aromatic C-H bond (Caro-H) is shorter than that in the alkyl C-H bond (Calk-H) due to a higher polarity of Caro-H, whereas the nuclear positions hardly change between the two groups. We found little difference in peak distances between Calk-H and Caro-H in the cryo-EM map. These observations are explained by the dominance of nuclear charges in electron scattering.

**Table 1.** Distance between the parent atom and hydrogen density peak or nucleus or electron in the hydrogen atom.

| Bond type <sup>a</sup> | Distance in Å |  |  |  |
| --- | --- | --- | --- | --- |
|  | hydrogen peak |  | hydrogen nucleus <sup>b</sup> | hydrogen electron <sup>c</sup> |
| | $\geq 2\sigma$ | $\geq 4\sigma$ | | |
| C-H <sup>d</sup> | 1.15 (0.13) <sup>g</sup><br>228 <sup>h</sup> | 1.09 (0.08)<br>43 | - | - |
| C<sub>alk</sub>-H <sup>e</sup> | 1.15 (0.12)<br>185 | 1.09 (0.09)<br>39 | 1.09 | 0.97 |
| C<sub>aro</sub>-H <sup>f</sup> | 1.14 (0.15)<br>43 | 1.10 (0.06)<br>4 | 1.08 | 0.93 |
| C-H<sub>2</sub> | 1.15 (0.12)<br>296 | 1.10 (0.11)<br>62 | 1.09 | 0.97 |
| C-H<sub>3</sub> | 1.17 (0.09)<br>181 | 1.09 (0.06)<br>28 | 1.09 | 0.97 |
| N-H <sup>i</sup> | 1.06 (0.09)<br>187 | 1.03 (0.07)<br>83 | 1.02 | 0.86 |
| O-H | 1.04 (0.07)<br>13 | 1.07 (0.10)<br>3 | 0.98 | 0.84 |

<sup>a</sup> Only for hydrogen densities detected in this structure excluding water.
<sup>b</sup> Derived by neutron crystallography and <sup>c</sup> by X-ray crystallography (34).
<sup>d</sup> Both the alkyl and aromatic groups, <sup>e</sup> the alkyl group and <sup>f</sup> the aromatic groups.
<sup>g</sup> The standard deviations in parentheses, and <sup>h</sup> number of observed peaks at the lower row of the cell.
<sup>i</sup> Peptide bonds, and side chains of asparagine, glutamine, histidine and arginine. Excluding lysine side chains due to facing to solvent.

The peak distances at density levels ≥ 2*σ* are longer by 0.1 – 0.3 Å than nuclear locations. We then plotted the hydrogen peak position vs B-factor and the peak position vs peak height for each bond type (Figs. S4 and S5). Here, B-factors along the horizontal axis of the plots are adopted from the parent non-hydrogen atom. Although distributions are noisy and numbers in Caro-H and O-H are small, there are consistent trends showing that peak locations become longer for higher B-factors and lower peak heights.

We finally estimated coordinate errors in the atomic model. In macromolecular X-ray crystallography, diffraction-component precision index (DPI) (32) is a standard measure for this. DPI relies on accuracy of diffraction intensity, but again CTF changes amplitude in EM images, yielding less accurate measures for amplitude. Still, REFMAC5 can provide DPI during refinement of a model against a single particle reconstruction, but we think a more suitable criterion is needed here. According to Cruickshank, the root mean square differences (RMSD) in atomic positions <Δ*r*> between two structures solved independently are in good agreement with the estimated standard uncertainties of positions *σ*(*r*) (32, 33). Accordingly, we calculated RMSD between two models built on two half maps reconstructed independently to estimate the standard uncertainty in atomic positions in the model. RMSD_1/2_, named here, and DPI obtained from REFMAC5 gave similar values, ~ 0.005Å RMSD_1/2_ could be used for estimation of coordinate errors in single particle cryo-EM models. For unrestrained refinement of crystal structures of small molecules, the standard uncertainty of a bond length between two atoms with similar coordinate errors is approximated by multiplying the coordinate error, i.e. DPI or *σ*(*r*), by the square root of 2 (32, 33), and this yields 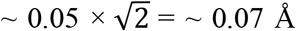. The standard deviations of the distances between the hydrogen peak density and the parent atom in Table 1 and Figs. S4 and S5 are comparable to 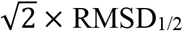. Considering that RMSD_1/2_ is calculated based on half maps and might be overestimated, differences between the average lengths of N-H and C-H_n_ bonds are 1 – 2 *σ* level (Table 1). Thus, the error level supports the observed dependence of the peak position on bonded-atom type.

## Conclusion

In this report, we describe detailed features of a high-resolution cryo-EM map and show that there are the same trends of the hydrogen density positions as observed in structures of an organic molecule revealed by 3D ED and X-ray free electron laser (XFEL) (27). Through the 3D ED and XFEL study, we noted that electrons could provide better hydrogen signals not only due to the relatively higher scattering power but also to the distal location of the density and to their higher sensitivity to charges. In addition to this, single particle analysis benefits substantially from accurate phase information derived directly from real images. Therefore, the sub-1.2 Å cryo-EM map yields better hydrogen signals than X-ray crystallography at this resolution range. Furthermore, selection of spatial frequencies for calculation of difference maps shows characteristic behaviors for charged atoms, which reveals negative charges on acidic-amino acids. Thus, this work demonstrates new potential of high-resolution single particle cryo-EM, and the technique could now become a powerful tool for analysis of detailed hydrogen properties and charges.

## Data availability

A cryo-EM density map and the atomic model of apoferritin for Dataset A have been deposited in the Electron Microscopy Data Bank and the Protein Data Bank (PDB) with accession codes EMD-32410 and 7WBY, respectively. Raw movies of this dataset have been deposited in the Electron Microscopy Public Image Archive with an accession code EMPIAR-ZZZZZ.

## Acknowledgements

We thank Sohei Motoki, Fumiaki Makino and Bartosz Marzec for technical support for axial-coma-free alignment and implementation of JAFIS Tool, Hisashi Naitow for setting up RELION, Yuko Kageyama for help in sample preparation, Kazuto Arakawa for information on Debye-Scherrer rings of Pt, Haruaki Yanagisawa for providing a plasmid of mouse apoferritin, and D. B. McIntosh for help in improving the manuscript. This work was partly supported by JST-Mirai Program Grant Number JPMJMI20G5 (to K. Y.) and the Cyclic Innovation for Clinical Empowerment (CiCLE) from the Japan Agency for Medical Research and Development, AMED (to K.Y.).

## Supplementary materials

### Materials & Methods

#### Sample preparation

Apoferritin was expressed in *E. coli* BL21-Gold(DE3) (Agilent) from a plasmid encoding mouse heavy-chain ferritin provided by Dr. Haruaki Yanagisawa, the University of Tokyo. The protein was purified by heat treatment at 70°C for 10 min, precipitation with ammonium sulfate, followed by gel filtration as described in the attached document. The apoferritin complex of ~ 500 kDa was eluted in 20 mM HEPES pH 7.5 and 300 mM NaCl, and the protein concentration was measured with the BCA assay (Pierce). Three μl of sample solution containing apoferritin at a concentration of 3 or 4 mg /ml was applied onto a holey carbon-coated grid (Quantifoil R1.2/1.3, Quantifoil Micro Tools GmbH) with 200 mesh, and the grid was blotted off with filter paper for 4 s and immediately plunge-frozen in liquid ethane using an FEI Vitrobot Mark IV (ThermoFisher Scientific) under 100% humidity at 4°C.

#### Adjustment of compensation for beam tilts and shifts

The spot size, illumination angle α and magnification were adjusted for the data acquisition condition. The object lenses were reset to the standard focus and the sample height (z) was brought to the just-focus position. The beam-tilt compensator (condenser lens deflector tilt; CL Comp Tilt) was first adjusted as per the manual provided by JEOL. Next, the illumination was set to the parallel condition, and an object was centred and defocused to −10 *μ*m with no condenser aperture inserted. Then, x and y beam shifts were changed by positive or negative adjustments a few μm at a time, and the overall image movement minimized with the beam-shift compensator (condenser lens deflector shift; CL Comp Shift). ChkLensDef in ParallEM (1) can store and restore CL Comp Tilt and Shift for each illumination angle α (4).

#### Data collection

The samples were examined at a specimen temperature of ~ 96 K with a CRYO ARM 300 electron microscope (JEOL) equipped with a cold-field emission gun and operated at an accelerating voltage of 300 kV. A parallel electron beam illuminated the sample, and inelastically scattered electrons were removed through an in-column energy filter with an energy slit width of 20 eV. Dose-fractionated images were recorded on a K3 camera (Gatan, AMETEK) in super-resolution mode with a nominal magnification of 100,000×, which corresponded to a physical pixel size of 0.495 Å.

All image data were acquired using SerialEM (18), and two datasets, Datasets A and B, were collected with and without JAFIS Tool ver. 1, respectively. JAFIS Tool is a python script developed by Dr. Bartosz Marzec, JEOL and can be called from the SerialEM script. It synchronizes image shifts with beam tilts and objective stigmas for removal of axial coma aberrations and two-fold astigmatism based on calibration, which can be prepared through a user-friendly GUI tool. The axial coma-free alignment was done as above for Dataset A before setting up JAFIS Tool. The Multiple Record setup and coma-free calibration in SerialEM was used for Dataset B.

Once the stage was aligned to a new hole by cross-correlation with the reference hole image, image shifts were applied over neighbouring holes around the central hole. One image stack was recorded from each of 5 × 5 holes (Dataset A) and 3 × 3 holes (Dataset B). The settings of lenses and deflector coils were monitored with ChkLensDef in ParallEM (1, 4).

In Dataset A, a total of 12,114 image stacks was collected in CDS mode at a dose rate setting of 3.819 e^-^·/ s per physical pixel and 0.0585 s per frame with total 2.34 s exposure and a defocus setting range of −0.5 to −1.0 μm. In Dataset B, a total of 2173 was in non-CDS mode at a dose rate setting of 15.872 e^-^·/ s per physical pixel and 0.016 s per frame with total 0.8 s exposure and a defocus setting range of −0.5 to −1.5 μm. The interval for flashing the gun was set to 8 h. Imaging parameters are shown in Table S1.

#### Structure analysis

We first removed micrographs taken from the regions less than 1.5 μm apart from those exposed before. This had resulted from stage alignment errors. A python script named distpos.py was made to find those data based on x, y coordinates in SerialEM meta data files (.mdoc). Subsequent image processing was carried out with RELION-3.1 (12, 35). Image stacks were drift-corrected, dose-weighted, and summed with the MotionCor2 algorithm (36) implemented in RELION. Contrast transfer function (CTF) parameters were estimated with CTFFIND (37) and images showing poor Thon rings were removed by eye. A 3D reconstruction of apoferritin previously obtained (4) was low-pass filtered to 20 Å and used as templates for automatic particle picking. Picked-up particles were extracted in 120 × 120 pixel boxes with a pixel size of 1.98 Å and reference-free 2D classification were repeated twice to select ones in good class averages. In total 2,235,864 and 311,583 particles were selected and extracted in 480 × 480 pixel boxes with a pixel size of 0.495 Å from 7852 and 1122 micrographs for Dataset A and B, respectively, and applied to 3D auto-refinement with octahedral symmetry enforcement. Anisotropic magnification, beam tilts, and CTF parameters were refined for each micrograph, followed by estimation and correction of particle-based motions. The refinement steps were iterated a few times, and 3D maps were reconstructed with correction of the Ewald-sphere curvature. The resolutions of the maps were estimated to be 1.21 Å and 1.49 Å for Dataset A and B, respectively, by the standard post-processing procedure of RELION. Only particles in Dataset A were re-extracted in 600 × 600 pixel boxes with a pixel size of 0.396 Å and the same refinement steps were repeated. Reconstruction with Ewald-sphere correction gave the final map at 1.19 Å resolution. Removal of micrographs with refined scale factors (rlnGroupScaleCorrection) < 0.5 and x, y beam tilt deviations > 0.15 mrad from the averages yielded 2,104,187 particles, but this improved the resolution of the reconstructed map only by the order of 10^-3^ Å. The Rosenthal-Henderson plots (21) were calculated for both datasets by dividing the entire data into subgroups with various particle numbers. Ewald-sphere correction was applied to subgroups consisting of ≥ 3200 particles.

An atomic model of mouse apoferritin (PDB ID: 7KOD) was fitted onto the 1.19 Å resolution map and inspected using UCSF Chimera (38). The model was refined with ISOLDE (39) and REFMAC5 (40) without hydrogen atoms. Then, unfiltered cryo-EM half maps were cut out into a box of 320 × 320 × 320 pixels. A Fourier difference map was calculated between the cut-out maps and the model omitting hydrogen atoms by *Servalcat* (5) in CCP-EM. Hydrogen atoms and water molecules with a density level of ≥ 2*σ* and ≥ 4*σ* were picked out using PEAKMAX from the CCP4 suite (41). Hydrogen peaks were manually edited by referring to the riding positions in the protein model. Structure refinement was carried out while retaining hydrogen atoms associated with the parent atoms with an average B-factor ≤ 20 Å^2^, and removing all hydrogen atoms belonging to water molecules or amino acid residues showing multiple conformations. The hydrogen positions were fixed during refinement. Anisotropic and isotropic temperature factors were used for non-hydrogen atoms and hydrogen atoms, respectively. Refinement statistics of the model were calculated using the CCP-EM validation program (42) and are summarized in Table S1. Q-score was calculated by replacing disordered side chains to alanine (43). Structure figures were prepared with PyMOL (The PyMOL Molecular Graphics System, Schrödinger, LLC) and UCSF Chimera.

---

**Figure S1.**
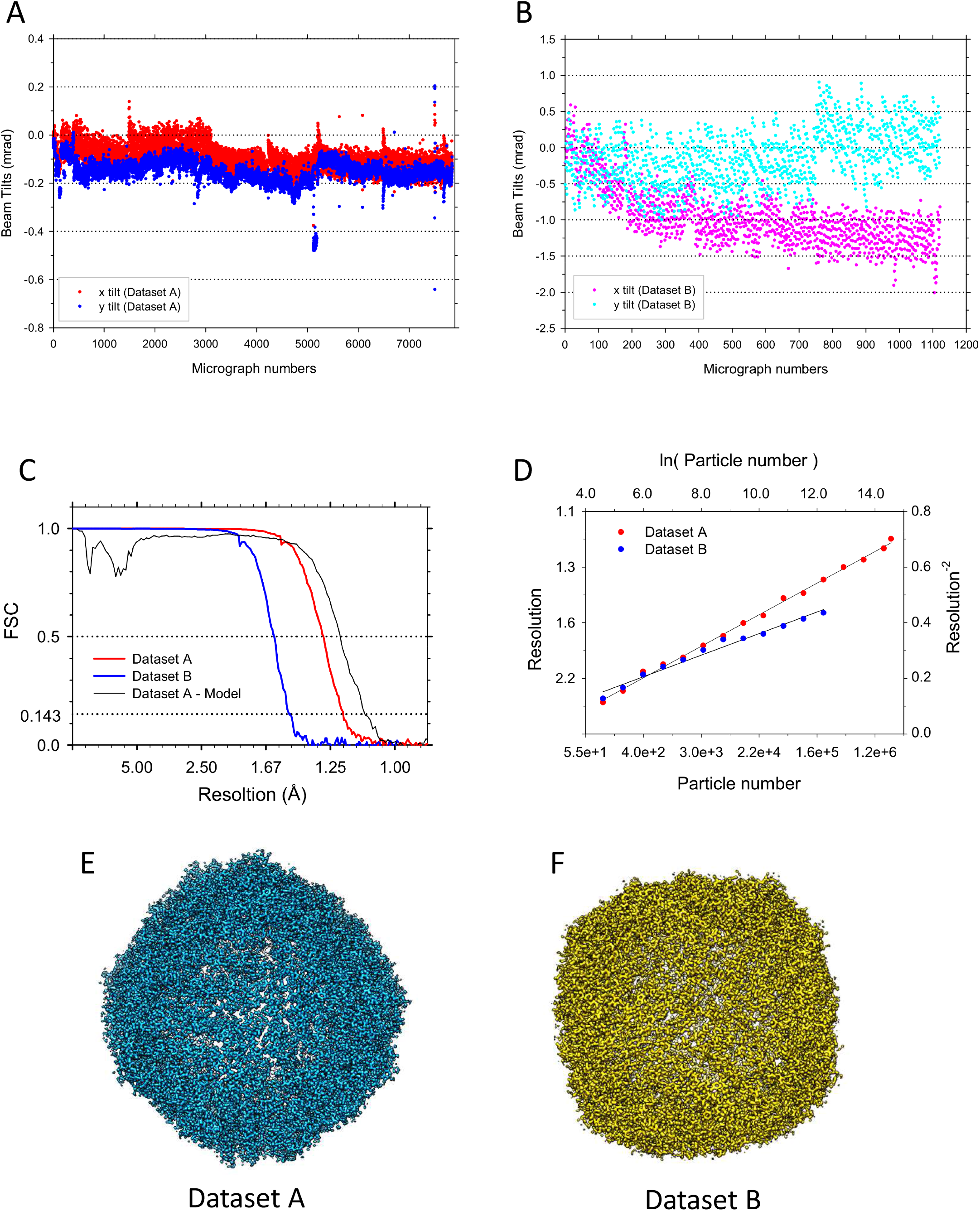
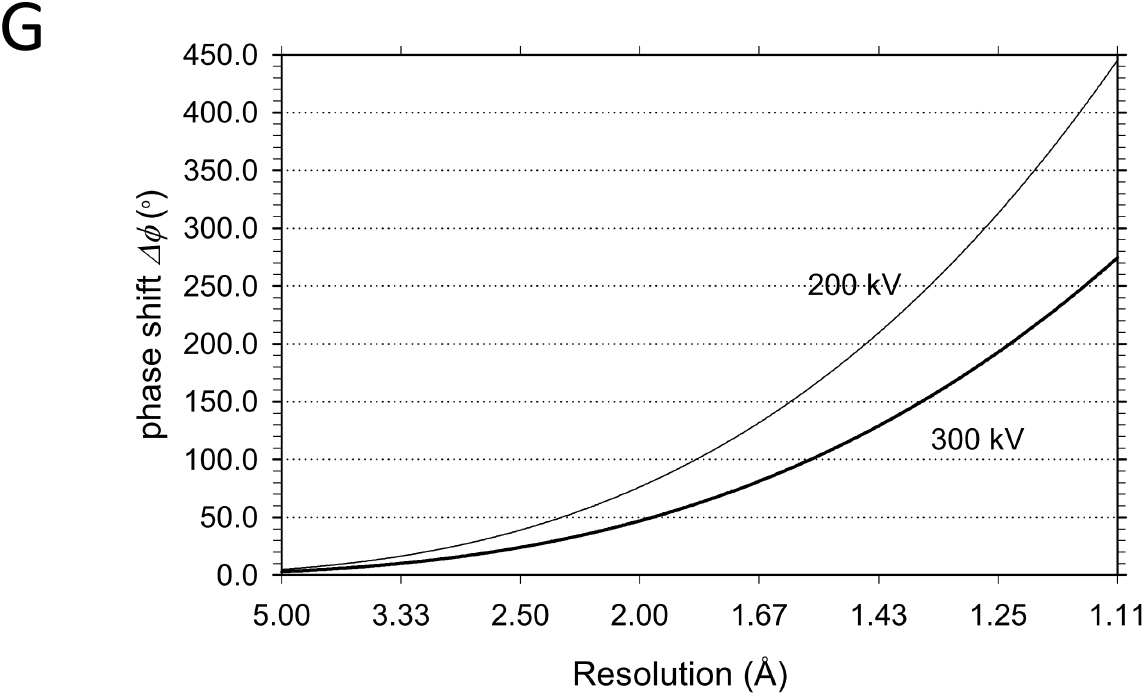
Comparison of Datasets A and B. (A) Plot of beam tilt variations for Dataset A. (B) Plot of beam tilt variations for Dataset B. (C) FSC plots for half maps of Datasets A, B and Dataset A vs the model. (D) Rosenthal-Henderson plots for Datasets A and B. The Rosenthal-Henderson B-factors are estimated as 34 Å^2^ for Dataset A and 51 Å^2^ for Dataset B. Reconstructions were repeated four times from subgroups consisting of ≤ 1600 randomly-selected particles, and Ewald-sphere correction was applied to subgroups of ≥ 6400 particles. (E) Reconstruction of Dataset A. (F) Reconstruction of Dataset B. (G) Plots of phase shift

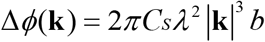

introduced by an angle off the axial coma-free axis *b* = 0.1 mrad at a spatial frequency **k** with a spherical aberration coefficient *Cs* = 2.7 mm and the wavelength of electron *λ* for 200 and 300 kV.

**Figure S2.**
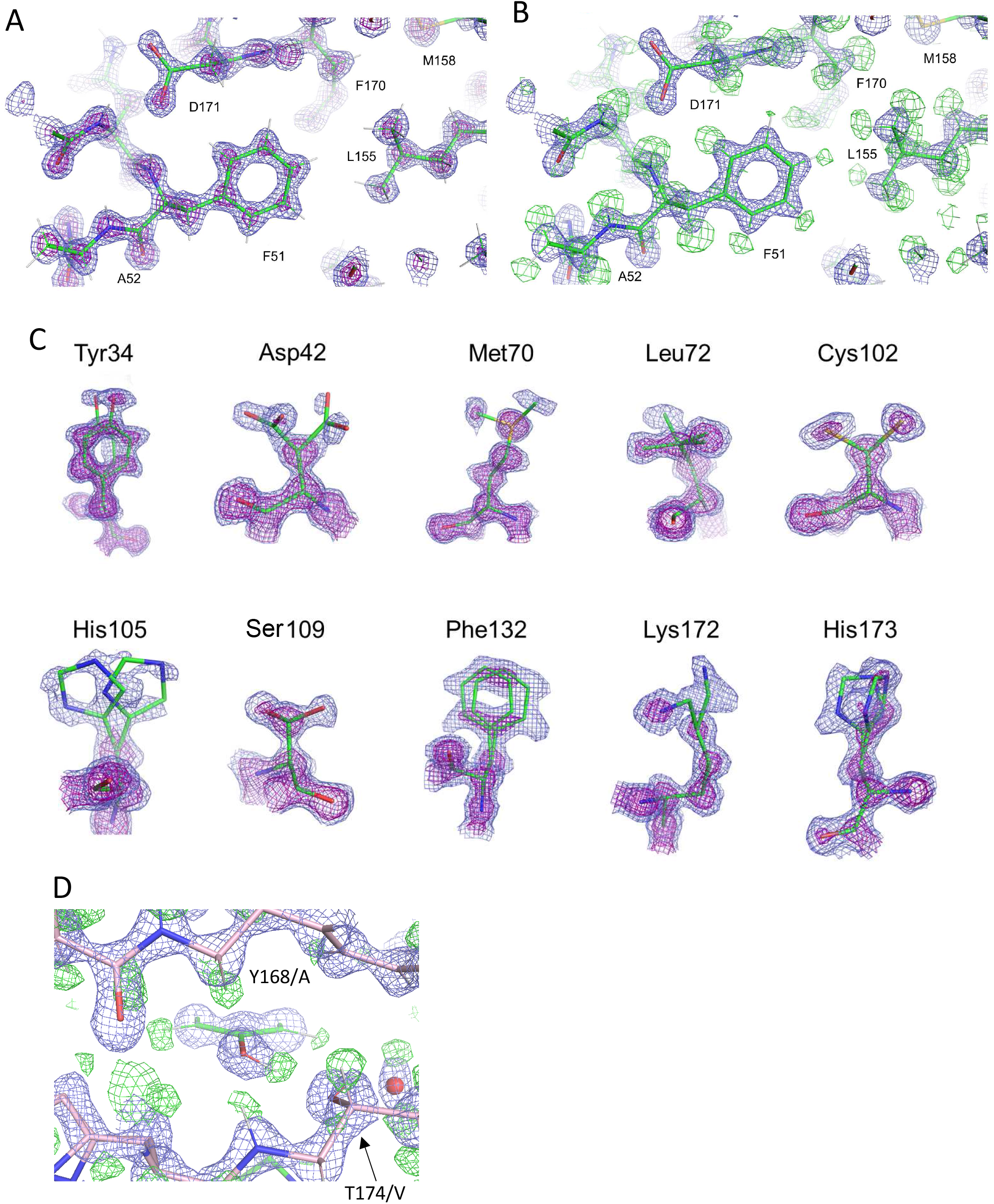
Other examples of structure details. (A, B) Around a phenylalanine (Phe 51). Displayed as in Figs. 2(A) and (B). (C) Amino-acid residues exhibiting multi-conformations. All the residues except for Tyr 34 are located on the protein surface exposed to solvent. The contour levels of blue and purple nets are 2*σ* and 7*σ*, respectively. (D) The same as in Fig. 2B but viewed roughly parallel to the phenyl ring from the hydroxyl group. The hydrogen atom (HH) at the hydroxyl group of Tyr 168 in subunit A forms a hydrogen bond with the side chain oxygen (OG) of Thr 174 in subunit V.

**Figure S3.**
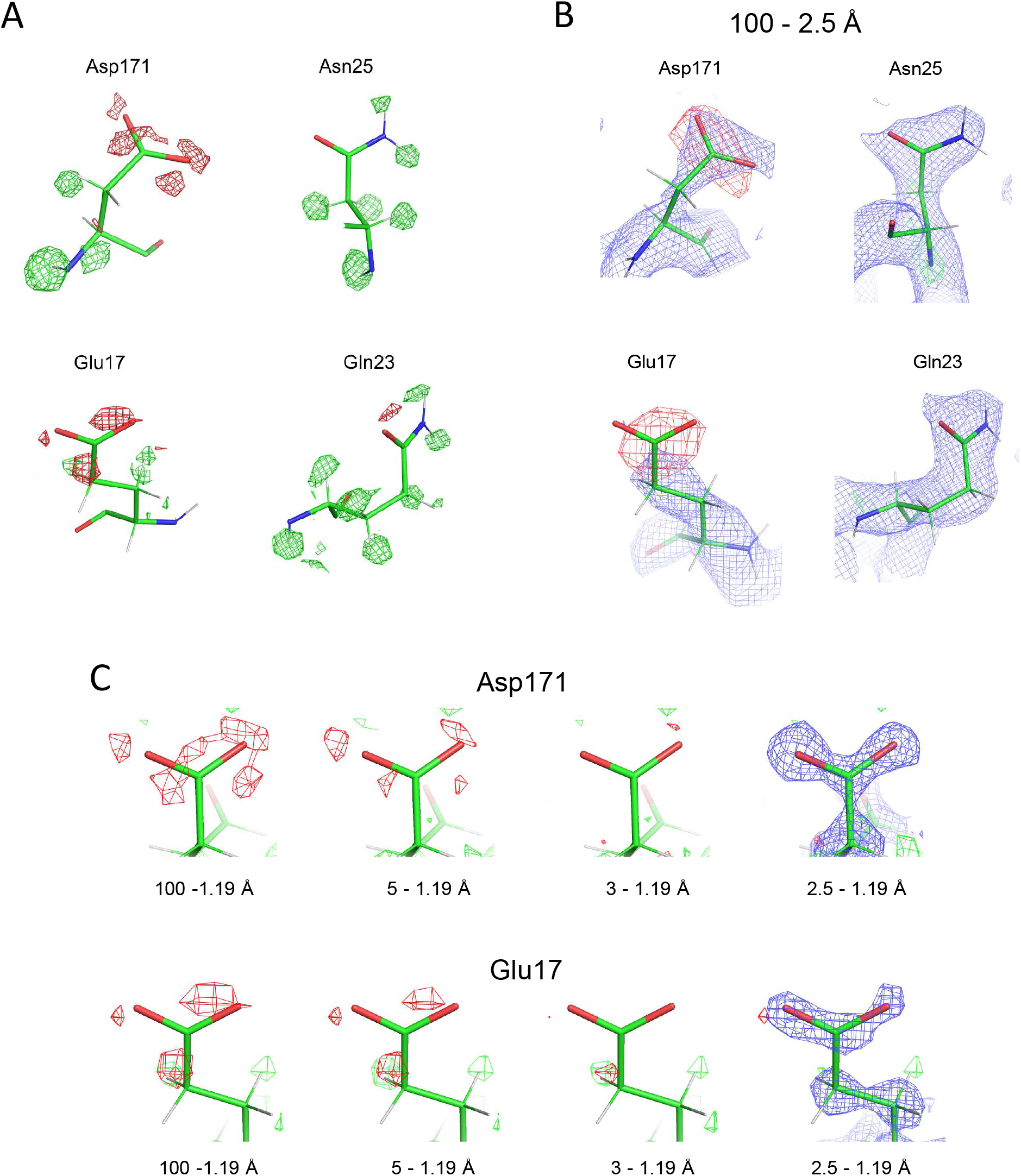
Analysis of negative densities. (A) The same as in Fig. 2F but overlaid with positive and negative densities in the difference map. B-factors for the terminal oxygen and bonded carbon atoms are: 23.3 Å^2^ (OD1), 20.2 Å^2^ (OD2) and 16.3 Å^2^ (CG) in Asp 171; 34.0 Å^2^ (OE1), 31.9 Å^2^ (OE2) and 21.7 Å^2^ (CD) in Glu 17; 19.2 Å^2^ (OD1) and 13.2 Å^2^ (CG) in Asn 25; and 24.6 Å^2^ (OE1) and 15.8 Å^2^ (CD) in Gln 23. (B) The same as in Fig. 2G (excluding the data in higher than 2.5 Å resolutions) but with the experimental map displayed in blue and at 2*σ*. The experimental map densities disappear for the terminal oxygen atoms of Glu 171. (C) Difference densities around side chains of Asp 171 and Glu 17. Calculated from inclusion of data in given resolution ranges. Green and red nets correspond to difference densities at levels of 4*σ* and −4*σ*, respectively. Experimental densities in blue are overlaid at 2*σ* only for those calculated from 2.5 – 1.19 Å resolution data.

**Figure S4.**
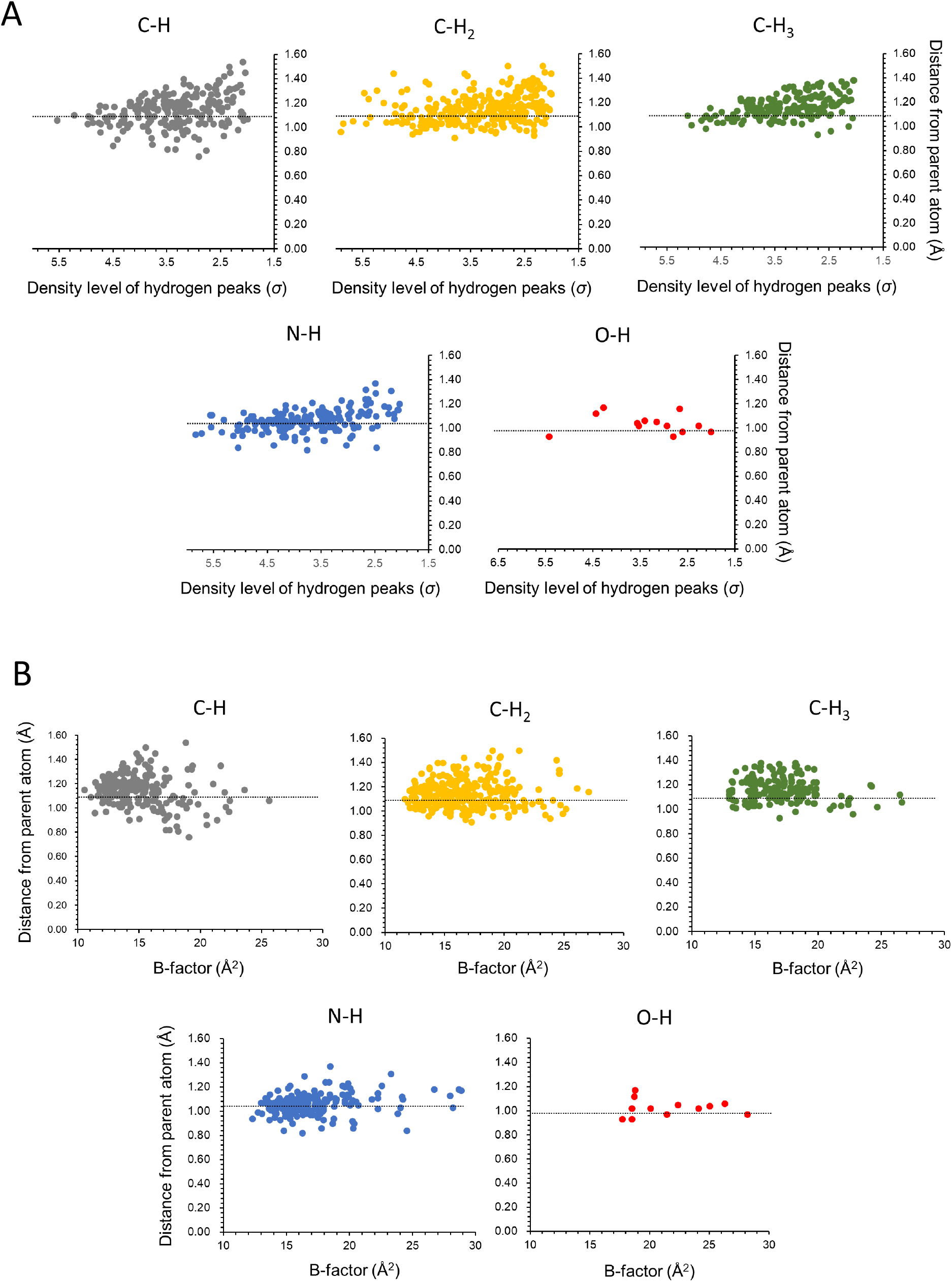
Plots of peak distance vs display level (A) and peak distance vs B-factor (B) for CH, C-H2, C-H3, N-H and O-H bonds. (C)

**Figure S5.**
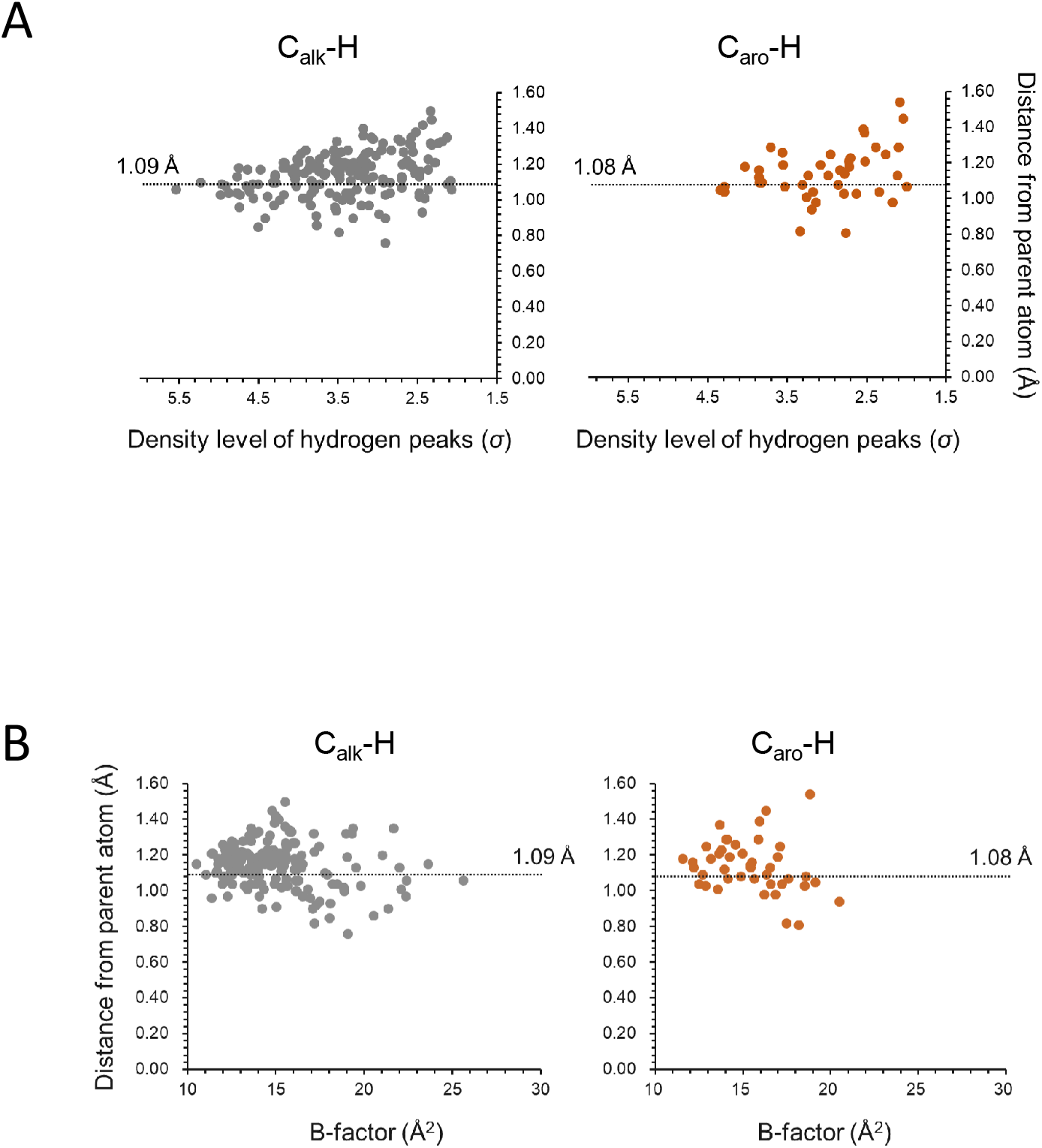
Plots of peak distance vs display level (A) and peak distance vs B-factor (B) for C_alk_-H and C_aro_- H bonds.

**Table S1.** Data and refinement statistics.

|  | Dataset A | Dataset B |
| --- | --- | --- |
| PDB-code | 7WBY | - |
| EMD-code | EMD-32410 | - |
| **Data collection** |  |  |
| Microscope | CRYO ARM 300 |  |
| Imaging device | K3 |  |
| Acceleration voltage (kV) | 300 |  |
| Imaging mode | CDS<br>Super-resolution | Counted<br>Super-resolution |
| Data collection | SerialEM + JAFIS Tool | SerialEM |
| Grid condition | Quantifoil R1.2/1.3 |  |
| Nominal magnification | x100,000 |  |
| Total exposure time (sec) | 2.34 | 0.8 |
| No. of frames | 40 | 50 |
| Total electron exposure (e<sup>-</sup>/Å<sup>2</sup>) | 36.47 | 51.82 |
| Defocus range (μm) | -0.5 - -1.0 | -0.5 - -1.0 |
| Physical pixel size (Å) | 0.495 | 0.495 |
| No. of total image sets | 12,114 | 2,173 |
| **Data processing** |  |  |
| No. of used image sets | 7,852 | 1,122 |
| Final particles (no.) | 2,235,864 | 311,583 |
| Pixel size for final map | 0.396 | 0.495 |
| Symmetry imposed | *O* | *O* |
| Map resolution (Å) | 1.19 | 1.49 |
| FSC threshold | 0.143 | 0.143 |
| **Refinement** |  |  |
| **Model composition in the asymmetric unit** |  |  |
| Initial models (PDB code) | 7KOD | - |
| Model resolution (Å) | 1.2 | - |
| FSC threshold | 0.5 | - |
| Q-score for protein * | 0.91 | - |
| Estimated resolution (Å) by Q-score for protein * | 1.18 | - |
| Model composition |  |  |
| Non-hydrogen atoms | 2248 | - |
| Protein residues | 172 | - |
| Water | 142 | - |
| Ion | 1 (Na<sup>+</sup>) | - |
| B-factors (Å<sup>2</sup>) |  |  |
| Protein | 19.5 | - |
| Water | 37.7 | - |
| Ion | 27.7 | - |
| R.m.s. deviations |  |  |
| Bond lengths (Å) | 0.013 | - |
| Bond angles (Å) | 1.55 | - |
| Validation |  |  |
| MolProbity score <sup>†</sup> | 1.09 | - |
| Clashscore | 3.00 | - |
| Poor rotamers (%) | 0.75 | - |
| Ramachandran plot |  |  |
| Favored (%) | 99.41 | - |
| Allowed (%) | 0.59 | - |
| Disallowed (%) | 0.00 | - |
| RMSD<sub>1/2</sub> (Å) <sup>‡</sup> | 0.05 | - |

\* An index showing resolvability of atoms, amino acid residues, and ligands assigned in a cryo-EM map (43).
<sup>†</sup> Defined in Ref. (34).
<sup>‡</sup> Defined in this work.

